# A genetic toolkit for studying transposon control in the *Drosophila melanogaster* ovary

**DOI:** 10.1101/2021.06.29.450424

**Authors:** Mostafa F. ElMaghraby, Laszlo Tirian, Kirsten-André Senti, Katharina Meixner, Julius Brennecke

## Abstract

Argonaute proteins of the PIWI class complexed with PIWI-interacting RNAs (piRNAs) protect the animal germline genome by silencing transposable elements. One of the leading experimental systems for studying piRNA biology is the *Drosophila melanogaster* ovary. In addition to classical mutagenesis, transgenic RNA interference (RNAi), which enables tissue-specific silencing of gene expression, plays a central role in piRNA research. Here, we establish a versatile toolkit focused on piRNA biology that combines germline transgenic RNAi, GFP marker lines for key proteins of the piRNA pathway, and reporter transgenes to establish genetic hierarchies. We compare constitutive, pan-germline RNAi with an equally potent transgenic RNAi system that is activated only after germ cell cyst formation. Stage-specific RNAi allows us to investigate the role of genes essential for germline cell survival, for example nuclear RNA export or the SUMOylation pathway, in piRNA-dependent and independent transposon silencing. Our work forms the basis for an expandable genetic toolkit provided by the Vienna Drosophila Resource Center.

## INTRODUCTION

Transposable elements (TEs) are mobile, selfish genetic elements that have parasitized almost all eukaryotic genomes and pose a threat to genome integrity (Feschotte 2008; Fedoroff 2012). In plants, fungi, and animals, small RNA silencing pathways are centrally involved in TE silencing, indicating an ancient function of RNA interference pathways in protecting the genome (Malone and Hannon 2009). In the animal germline, genome defense guided by small RNAs is carried out by Argonaute proteins of the PIWI-clade and their bound PIWI-interacting RNAs (piRNAs) (Siomi *et al*. 2011; Czech *et al*. 2018; Ozata *et al*. 2018). Most piRNAs originate from discrete genomic loci called piRNA clusters, which are rich in TE insertions (Aravin *et al*. 2007; Brennecke *et al*. 2007; Houwing *et al*. 2007). Therefore, piRNAs can guide PIWI proteins to complementary TE transcripts, allowing their selective silencing at the transcriptional (nuclear PIWIs) and post transcriptional (cytoplasmic PIWIs) levels. Defects in the piRNA pathway are compatible with overall animal development but result in uncontrolled TE activity in gonads, DNA damage, ectopic recombination and sterility. As stable cell lines derived from germline cells are rare, and as the piRNA pathway can differ in different cell types, the arms race between TEs and the piRNA pathway must be studied within the context of a developing organism.

*Drosophila* oogenesis is one of the leading model systems for piRNA research. Two main cell types make up the fly ovary: (1) germline cells (germline stem cells, dividing cystoblast cells, nurse cells and oocyte) and (2) somatic support cells that form the stem cell niche and surround, nourish, and protect the germline cells (Hudson and Cooley 2014). Both, germline and somatic cells of the ovary harbor a functional piRNA pathway. However, these pathways differ in several aspects. For example, germline cells express three PIWI clade Argonautes (nuclear Piwi, cytoplasmic Aubergine and Ago3), whereas somatic cells of the ovary express only nuclear Piwi (Malone *et al*. 2009). The identity and biology of the genomic piRNA source loci also differ in the two cell types. piRNA clusters in the ovarian soma resemble canonical RNA Polymerase II transcription units with defined promoter, transcription start site and termination site (Lau *et al*. 2009; Malone *et al*. 2009; Goriaux *et al*. 2014; Mohn *et al*. 2014). Germline piRNA clusters are instead specified at the chromatin level by the action of Rhino, a member of the heterochromatin protein 1 (HP1) family that recruits germline specific variants of core gene expression factors to enable enhancer-independent transcription on both DNA strands within heterochromatic loci (Klattenhoff *et al*. 2009; Mohn *et al*. 2014; Zhang *et al*. 2014; Andersen *et al*. 2017). The resulting piRNA precursors are suppressed in splicing and canonical 3’ end formation and are exported via a dedicated, germline specific RNA export route to the cytoplasmic, perinuclear piRNA processing sites known as nuage (Chen *et al*. 2016; ElMaghraby *et al*. 2019; Kneuss *et al*. 2019).

Defects in the germline piRNA pathway result in uncontrolled TE transposition and persistent activation of the DNA damage checkpoint. As a result, oocyte patterning pathways are disrupted, and eggs laid by piRNA pathway mutant flies have dorso-ventral polarity defects (Theurkauf *et al*. 2006; Klattenhoff *et al*. 2007; Senti *et al*. 2015; Durdevic *et al*. 2018; Wang *et al*. 2018). Based on this phenotype, classic genetic screens have uncovered several piRNA pathway genes (Schupbach and Roth 1994; Cook *et al*. 2004; Wehr *et al*. 2006; Chen *et al*. 2007; Pane *et al*. 2007; Zamparini *et al*. 2011). With the development of transgenic RNAi and the establishment of genome-wide *Drosophila* RNAi libraries (Dietzl *et al*. 2007; Haley *et al*. 2008; Ni *et al*. 2008; Ni *et al*. 2011), reverse genetic screens have systematically revealed piRNA pathway genes (Czech *et al*. 2013; Handler *et al*. 2013). Depending on the Gal4 driver used, these screens were specific to the somatic or germline piRNA pathway. Together, they identified ∼40 genes with specific functions in the piRNA pathway.

Transgenic RNAi in the germline is based on *nanos*-Gal4 driver lines that activate the expression of long or short RNA hairpin constructs targeting a gene of interest. The two most powerful transgenic RNAi setups for the germline are: (1) Combining the Maternal Triple Driver (MTD-Gal4; a combination of COG-Gal4 on the X-chromosome, NGT-Gal4 on the second, and *nanos*-Gal4 on the third chromosome; (Grieder *et al*. 2000) with transgenes harboring short hairpins (shRNA; microRNA mimics) under UAS-control (Valium20/22 backbones from the Harvard TRiP collection) (Ni *et al*. 2011). And (2), the combination of a dual nanos-Gal4 driver line that activates the expression of UAS-controlled long dsRNA hairpins from the Vienna RNAi collection and of the RNAi-boosting protein Dcr-2 (Dietzl *et al*. 2007; Wang and Elgin 2011). Both approaches result in potent silencing of target gene expression throughout oogenesis, from primordial germ cells to germline stem cells to nurse cells and mature oocytes.

While the pan-germline knockdown approaches have been instrumental for piRNA pathway research, they are not without limitations. For example, the piRNA pathway intersects with several general cellular processes such as SUMOylation, transcription, chromatin modification, and RNA export. Genetic disruption of these processes often leads to cell-lethal or pleiotropic phenotypes resulting in atrophic ovaries lacking germline cells, precluding meaningful analysis. Previous studies have identified and characterized alternative Gal4 driver lines that activate Gal4 expression in the female germline upon cystoblast differentiation (late germarium stages onward), leaving germline stem cells unaffected (Staller *et al*. 2013). This allows genes with cell-essential functions to be studied as ovarian development proceeds to a sufficient extent.

Here, we first combine pan-oogenesis transgenic RNAi with marker lines expressing GFP-tagged piRNA pathway proteins with diverse molecular functions and sub-cellular localization. This toolkit provides a cell biology assay system for studying gene function within the germline piRNA pathway. We then introduce and characterize TOsk-Gal4, which causes strong Gal4 expression in the female germline immediately after germline cyst formation. TOsk-Gal4 is compatible with short and long hairpin RNAi libraries and allows efficient depletion of genes essential for cell survival without drastically affecting ovarian morphology and integrity. We combine TOsk-Gal4 with various genetic and molecular tools to study the interface between piRNA pathway, RNA export and protein SUMOylation.

## MATERIALS & METHODS

### Fly Husbandry

Flies were maintained at 25°C under light/dark cycles and 60% humidity. For ovary dissection, flies were kept in cages on apple juice plates with yeast paste for at least 5 days before dissection. All fly strains used and generated in this study are listed in Supplementary Table 1 and available via VDRC (https://stockcenter.vdrc.at/control/main).

### Generation of transgenic fly strains

We generated fly strains harboring short hairpin RNA (shRNA) expression cassettes by cloning shRNA sequences into the Valium-20 vector (Ni *et al*. 2011) modified with a white selection marker. Tagged fly stains were generated via insertion of desired tag sequences into locus-containing Pacman clones (Venken *et al*. 2009) via bacterial recombineering (Ejsmont *et al*. 2011).

### RNA Fluorescent In Situ Hybridization (RNA-FISH)

5-10 ovary pairs were fixed in IF Fixing Buffer for 20 minutes at room temperature, washed three times for 10 minutes in PBX, and permeabilized in 70% ethanol at 4 °C overnight. Permeabilized ovaries were rehydrated in RNA-FISH wash buffer (10% (v/w) formamide in 2× SSC) for 10 minutes. Ovaries were resuspended in 50 µl hybridization buffer (10% (v/w) dextran sulfate, 10% (v/w) formamide in 2× SSC) supplemented with 0.5 µl of 25 µM RNA-FISH probe set solution (Stellaris; Supplementary Table 2 lists oligo sequences). Hybridization was performed at 37 °C overnight with rotation. Next, ovaries were washed twice with RNA-FISH wash buffer for 30 minutes at 37 °C, and twice with 2xSSC solution for 10 minutes at room temperature. To visualize DNA, DAPI (1:10,000 dilution) was included in the first 2xSSC wash. Ovaries were mounted in ∼40 µl Prolong Diamond mounting medium and imaged on a Zeiss LSM-880 confocal-microscope with AiryScan detector.

### Immunofluorescence staining of ovaries

5-10 ovary pairs were dissected into ice-cold PBS and fixed in IF Fixing Buffer (4 % formaldehyde, 0.3 % Triton X-100, 1x PBS) for 20 minutes at room temperature with rotation. Fixed ovaries were washed thrice with PBX (0.3 % Triton X-100, 1x PBS), 10 minutes each wash, and incubated in BBX (0.1% BSA, 0.3 % Triton X-100, 1x PBS) for 30 minutes for blocking. Primary antibodies diluted in BBX were added to ovaries and binding was performed at 4°C overnight. After three 10 minute-washes in PBX, ovaries were incubated with secondary antibodies (1:1000 dilution in BBX) at 4°C overnight. Afterwards, the ovaries were washed 4 times with PBX, with the second wash done with DAPI (1:50,000 dilution). To visualize the nuclear envelope, Alexa Fluor 647-conjugated wheat germ agglutinin (1:200 dilution in PBX; Thermo Fisher Scientific) was added after DAPI staining for 20 minutes. Ovaries were finally mounted in ∼40 µl Prolong Diamond mounting medium and imaged on a Zeiss LSM-880 confocal-microscope with AiryScan detector. The resulting images processed using FIJI/ImageJ (Schindelin *et al*. 2012). Supplementary Table 3 list antibodies used in this study.

### Western blot analysis

10 ovary pairs were dissected in ice-cold PBS and homogenized with a plastic pestle in RIPA lysis buffer (50 mM Tris-HCl pH 7.5, 150 mM NaCl, 1% Triton X-100, 0.5% Na-deoxycholate,, 0.1% SDS, 0.5 mM EGTA, 1 mM EDTA) freshly supplemented with 1mM Pefabloc, cOmplete Protease Inhibitor Cocktail (Roche), and 1 mM DTT. The samples were spun down at 14,000 rpm for 5 minutes, and the homogenization step was repeated. After pooling both supernatants, samples were incubated on ice for 30 minutes and cleared by centrifugation at 14,000 rpm for 15 minutes. Protein concentrations were quantified by Bradford reagent, and 10 µg protein were resolved by SDS-polyacrylamide gel electrophoresis and transferred to a 0.2 µm nitrocellulose membrane (Bio-Rad). The membrane was blocked in 5% skimmed milk powder in PBX (0.01% Triton X-100 in PBS) and incubated with primary antibody overnight at 4°C (Supplementary Table 3). After three washes with PBX, the membrane was incubated with HRP-conjugated secondary antibody for 1h at room temperature, followed by three washes with PBX. Subsequently, the membrane was covered with Clarity Western ECL Blotting Substrate (Bio-Rad) and imaged using the ChemiDoc MP imaging system (Bio-Rad).

### RNA-Seq library preparation

5 ovary pairs were homogenized with a plastic pestle in 200 µL TRIzol reagent, and after homogenization 800 µL TRIzol were added and incubated for 5 minutes at room temperature. 200 µL chloroform–isoamyl-alcohol (24:1; Sigma Aldrich) were added, and after vigorous shaking, samples were incubated for 5 minutes at room temperature. Next, samples were centrifuged at 12,000 g for 15 minutes at 4°C. RNA was transferred from the upper aqueous phase using the Direct-zol RNA Miniprep kit (Zymo Research) with in-column DNaseI treatment according to manufacturer’s instructions. rRNA depletion from 1 µg total RNA was performed as described previously (ElMaghraby et al., 2019). Libraries were then cloned using the NEBNext Ultra II Directional RNA Library Prep Kit for Illumina (NEB), following the recommended kit protocol and sequenced on a NovaSeq 6000 – SR100 (Illumina).

### RT-qPCR analysis of transposon expression

Five pairs of dissected ovaries were homogenized in TRIzol reagent followed by RNA purification according to the manufacturer’s protocol. RNA was further purified with Direct-zol MiniPrep kit (Zymo Research) with DNase I treatment. 1 µg of total RNA was reverse transcribed using the Maxima First Strand cDNA Synthesis kit (Thermo Fischer) following standard protocols. cDNA was used as template for RT-qPCR quantification of transposon mRNA abundances (for primer sequences see Supplementary Table 4).

### Data availability

Table S1 list all fly stocks used and generated in the study. All fly stocks are available via the Vienna Drosophila Resource Center (VDRC). Next-Generation Sequencing data produced in this publication has been deposited to the NCBI GEO archive under the accession number GSE174611. Figure S1 indicates a crossing scheme for generating Gal4-based reporter lines. Table S2 lists of Stellaris RNA-FISH probes used in this study. Table S2 and S3 list antibodies and oligo sequences used respectively. The authors affirm that all data necessary for confirming the conclusions of the article are present within the article, figures, and tables.

## RESULTS AND DISCUSSION

### Combining pan-oogenesis RNAi with GFP-based piRNA pathway marker transgenes

Approximately forty proteins act in the *Drosophila* piRNA pathway. These factors serve different molecular functions and are localized to distinct subcellular locations in the nucleus (e.g. nucleoplasm or genomic piRNA source loci) and/or in the cytoplasm (e.g. outer mitochondrial membrane, perinuclear nuage). To visualize piRNA pathway proteins in whole mount ovary preparations by confocal microscopy, we generated transgenic fly lines carrying genomic rescue constructs with a FLAG-GFP tag at the N- or C-terminus of key piRNA pathway factors (four examples shown in Figure 1A). GFP-tagging allows accurate and semi-quantitative determination of the subcellular localization of a protein as it circumvents the limitations of antigen accessibility to primary and secondary antibodies. This is particularly relevant in late stage egg chambers (Figure 1B) or for factors enriched in peri-nuclear nuage such as Nxf3, Bootlegger, or Nibbler (Figure 1C).

**Figure 1:**
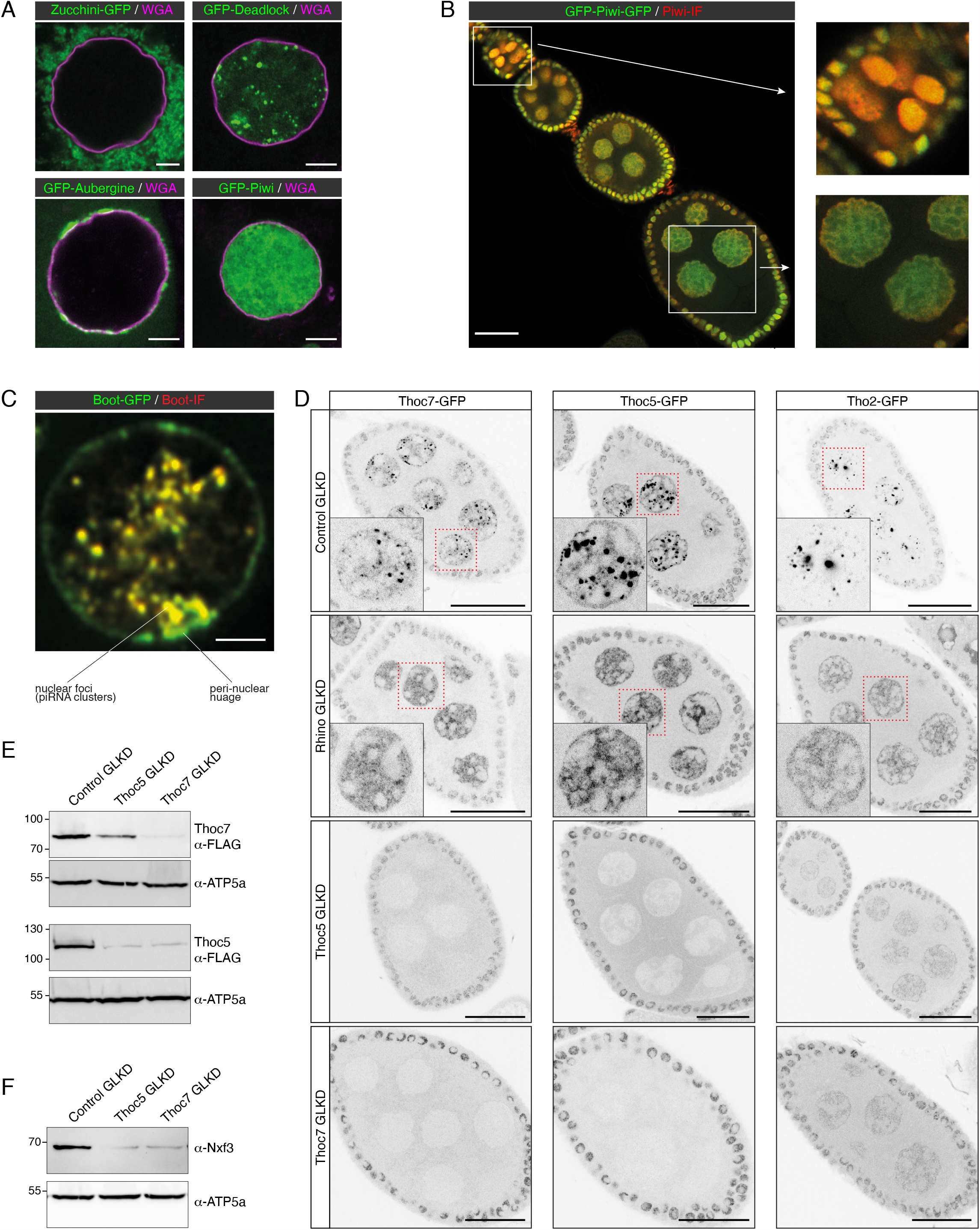
Pan-oogenesis RNAi with GFP-based piRNA pathway marker transgenes. (A) Confocal images (scale bars: 5 μm) showing localization of GFP-tagged Zucchini (mitochondrial membrane), Deadlock (piRNA clusters), Aubergine (cytoplasm with nuage enrichment), and Piwi (nuclear) in germline nurse cell nuclei (nuclear envelope labelled with WGA in magenta). (B) Confocal image (scale bar: 25 μm) showing GFP-Piwi localization (green) in an ovariole stained also with anti-Piwi antibody (red). To the right, an enlarged early egg chamber with good overlap between GFP and immunofluorescence (IF) signals (top) and nurse cell nuclei from an older egg chamber where the GFP signal dominates due to reduced antibody penetration (bottom) are shown. (C) Confocal image (scale bar: 3 μm) showing a nurse cell nucleus expressing Bootlegger-GFP (green) from the endogenous locus stained with an anti-Bootlegger antibody (red). Overlap between GFP and IF signals is apparent in nuclear foci, yet very poor in nuage. (D) Confocal images of egg chambers (scale bars: 25 μm) displaying localization of GFP-tagged Thoc7, Thoc5, or Tho2 (greyscale) in the indicated germline knockdown (GLKD) conditions (nuclei highlighted in red are enlarged). (E-F) Western blot analysis indicating levels of Thoc7 and Thoc5 (E), or Nxf3 (F) in ovary lysates from flies with indicated genotype (anti ATP-synthase blots served as loading control).

To be able to analyze the subcellular localization of the different piRNA pathway proteins in flies with targeted genetic perturbations (using transgenic RNAi), we combined the established GFP marker lines with germline specific Gal4 drivers. The resulting fly strains can be crossed with genome-wide collections of UAS lines that allow expression of long or short double stranded RNA constructs targeting any gene of interest (available from VDRC or Bloomington/TRiP). Figure S1 shows the crossing schemes underlying the construction of MTD-Gal4 lines (compatible with short hairpin RNA (shRNA) UAS-lines) or *nanos*-Gal4 lines with a UAS-Dcr2 transgene (compatible with long hairpin RNA UAS-lines) harboring the various GFP reporter transgenes. The resulting stocks represent a core set of piRNA marker lines that can be crossed with available RNAi stocks and that are available from the VDRC (Table 1).

**Table 1.** MTD-Gal4 reporter lines

|  | Name | Gene name | CG Number | Source |
| --- | --- | --- | --- | --- |
| THO/TREX | MTD-Gal4 + Thoc5-GFP | *thoc5* | CG2980 | This paper |
|  | MTD-Gal4 + GFP-Thoc7 | *thoc7* | CG17143 | This paper |
|  | MTD-Gal4 + GFP-Tho2 | *Tho2* | CG31671 | This paper |
|  | MTD-Gal4 + GFP-UAP56 | *Hel25E* | CG7269 | ElMaghraby *et al.*, 2019 |
| piRNA cluster biology | MTD-Gal4 + GFP-Rhino | *rhi* | CG10683 | Mohn *et al.*, 2014 |
|  | MTD-Gal4 + GFP-Deadlock | *cuff* | CG13190 | Mohn *et al.*, 2014 |
|  | MTD-Gal4 + GFP-Cutoff | *del* | CG9252 | Mohn *et al.*, 2014 |
|  | MTD-Gal4 + GFP-Moonshiner | *moon* | CG12721 | ElMaghraby *et al.*, 2019 |
|  | MTD-Gal4 + GFP-Nxf3 | *nxf3* | CG32135 | ElMaghraby *et al.*, 2019 |
|  | MTD-Gal4 + GFP-Bootlegger | *boot* | CG13741 | ElMaghraby *et al.*, 2019 |
| piRNA biogenesis | MTD-Gal4 + GFP-Zucchini | *zuc* | CG12314 | Hayashi *et al.*, 2016 |
|  | MTD-Gal4 + GFP-Gasz | *Gasz* | CG2183 | This paper |
|  | MTD-Gal4 + GFP-Vasa | *vas* | CG46283 | This paper |
|  | MTD-Gal4 + GFP-Aubergine | *spn-E* | CG3158 | This paper |
|  | MTD-Gal4 + GFP-Spindle E | *aub* | CG6137 | This paper |
|  | MTD-Gal4 + GFP-Nibbler | *Nbr* | CG9247 | This paper |
| piRNA-mediated silencing and heterochromatin | MTD-Gal4 + GFP-Nxf2 | *nxf2* | CG4118 | This paper |
|  | MTD-Gal4 + GFP-Gtsf1 | *arx* | CG3893 | This paper |
|  | MTD-Gal4 + GFP-Panoramix | *Panx* | CG9754 | This paper |
|  | MTD-Gal4 + GFP-Piwi | *piwi* | CG6122 | This paper |
|  | MTD-Gal4 + GFP-Sov | *sov* | CG14438 | This paper |
|  | MTD-Gal4 + GFP-HP1 | *Su(var)3-9* | CG43664 | This paper |
|  | MTD-Gal4 + GFP-Eggless | *egg* | CG12196 | This paper |
|  | MTD-Gal4 + silencing reporter |  |  | Sienski *et al.*, 2015 |

To illustrate the utility of the system, we focused on three subunits of the hexameric THO complex. THO is a key factor for nuclear mRNP quality control and, together with the RNA helicase UAP56 and the adaptor protein Aly/Ref1, functions as a central gatekeeper for nuclear mRNP export (Heath *et al*. 2016). In germline cells, THO binds piRNA precursors derived from heterochromatic piRNA clusters in addition to mRNAs (Zhang *et al*. 2012; Hur *et al*. 2016; Zhang *et al*. 2018). Based on its central role in nuclear mRNA export, THO is also thought to be required for the export of piRNA precursors. Consistent with this, THO localizes broadly in the nucleoplasm in all cells, but is additionally enriched in germline cells at genomic piRNA source loci that are specified by the HP1 family protein Rhino (Hur *et al*. 2016; Zhang *et al*. 2018).

We generated MTD-Gal4 lines expressing GFP-tagged THO subunits Tho2, Thoc5, or Thoc7, and crossed them with UAS-shRNA lines targeting *rhino, thoc5* or *thoc7*. As expected, depletion of Rhino resulted in loss of Tho2, Thoc5, and Thoc7 accumulation in discrete nuclear foci, indicating that THO localization to piRNA clusters depends on Rhino (Figure 1D) (Hur *et al*. 2016; Zhang *et al*. 2018). Loss of Thoc5 or Thoc7 revealed a strict co-dependence between both proteins for their stability, while Tho2 levels were only moderately affected in ovaries lacking Thoc5 or Thoc7 (Figures 1D-E). However, Tho2 localization to nuclear foci (piRNA clusters) strictly depended on Thoc5 and Thoc7 (Figure 1D). Consistent with a critical role of THO at piRNA clusters, flies lacking Thoc5 or Thoc7 in the germline, despite having morphologically normal ovaries, were sterile. Their sterility was presumably linked to defects in piRNA precursor export, supported by instability of Nxf3, the dedicated RNA export receptor for piRNA precursors (Figure 1F). In line with this, flies with strong hypomorphic *thoc5* or *thoc7* alleles are viable but show loss of piRNAs from Rhino-dependent clusters and are sterile (Hur *et al*. 2016; Zhang *et al*. 2018). In contrast, depletion of Tho2 resulted in rudimentary ovaries, suggesting that Tho2 is genetically more important for mRNA export than the Thoc5 and Thoc7 subunits. These results are of interest in light of structural and biochemical studies of the human THO-UAP56 complex: Whereas Tho2 is part of the THO core assembly (alongside Hpr1 and Tex), Thoc5 and Thoc7 form an extended coiled coil that is responsible for dimerization of the hexameric THO complex (Puhringer *et al*. 2020). Our results illustrate that the combination of transgenic RNAi with GFP-transgenes is a powerful system to study protein function in the ovarian piRNA pathway and more generally during oogenesis.

A clear limitation of the pan-oogenesis Gal4 driver system is that genes, whose depletion is incompatible with oogenesis (e.g. *tho2*), cannot be studied. Transgenic RNAi of genes with cell-essential functions results in rudimentary ovaries lacking detectable germline cells. Numerous genes (e.g. those involved in heterochromatin establishment, SUMOylation, nuclear RNA export) that are required for a functional piRNA pathway can therefore not be studied in this manner. Inspired by previous studies (Staller *et al*. 2013; Yan *et al*. 2014), we set out to characterize alternative germline-specific Gal4 driver lines that, in combination with UAS-RNAi lines, are compatible with the analysis of cell-essential genes in the context of the ovarian piRNA pathway.

### Efficient and specific transgenic RNAi in the differentiating female germline

Several germline specific genes are transcribed in differentiating cysts but not during embryonic, larval and pupal stages or in germline stem cells of the adult ovary. Gal4 driver lines exist for two of these genes: *oskar (osk)* and *alpha-Tubulin at 67C* (*αTub67C*) (Figure 2A) (Benton and St Johnston 2003; Telley *et al*. 2012). The *αTub67C*-Gal4 driver has been shown to induce efficient short-hairpin based RNAi in ovaries (Staller *et al*. 2013; Yan *et al*. 2014). We set out to systematically compare *osk*-Gal4 and *αTub67C*-Gal4 to the pan-oogenesis MTD-Gal4. We first crossed each driver line with a fly line carrying a UASp-H2A-GFP transgene. In addition, we also induced H2A-GFP expression with *traffic jam* (*tj*)-Gal4, a somatic driver that is active in all somatic support cells of the ovary(Tanentzapf *et al*. 2007; Olivieri *et al*. 2010). Western blot analysis indicated that H2A-GFP levels in ovary lysate were comparable (*osk*-Gal4) to, or even higher *(αTub67C*-Gal4), than those from the MTD-Gal4 crosses (Figure 2B). However, in contrast to the MTD-Gal4 crosses, no H2A-GFP was detectable in germline stem cells and early germline cysts in the germarium for the *osk*-Gal4 or *αTub67C*-Gal4 crosses (Figures 2A, C). We consistently observed that H2A-GFP expression initiated slightly earlier (germarium region 2b) for *osk*-Gal4 than for *αTub67C*-Gal4 (stage 2 egg chamber).

**Figure 2:**
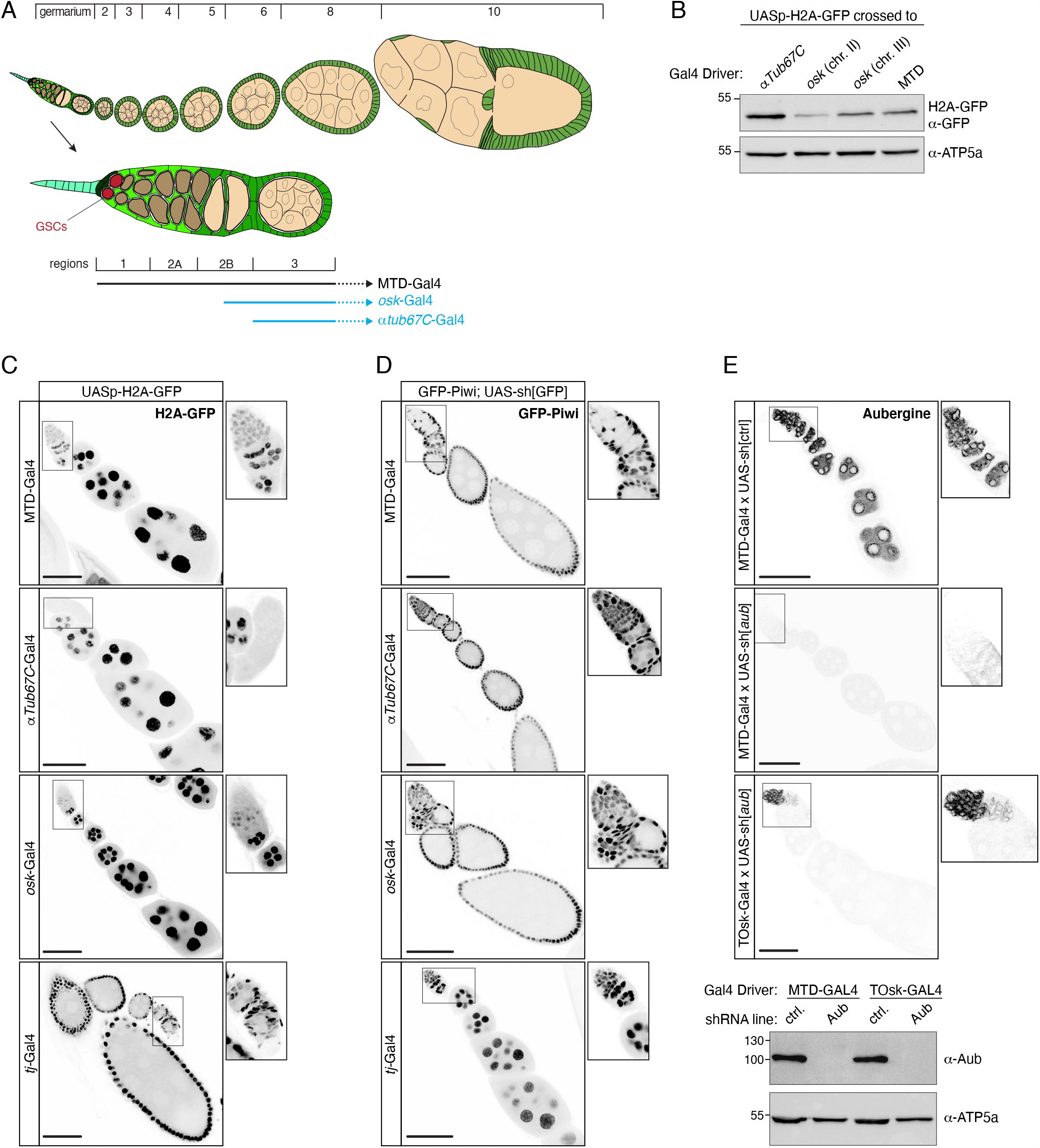
Efficient and specific transgenic RNAi in the differentiating female germline. (A) Cartoon of a *Drosophila* ovariole with somatic cells in green and germline cells in beige; egg chamber stages are indicated above and a magnified view of the germarium with stem cell niche is shown below. (B) Western blot analysis indicating levels of H2A-GFP expressed with indicated Gal4 drivers (anti ATP-synthase blot served as loading control). (C) Confocal images (scale bars: 50 μm) showing ovarioles expressing H2A-GFP (greyscale) driven by indicated germline and soma Gal4 drivers (captions to the right show enlarged germaria). (D) Confocal images (scale bars: 50 μm) showing ovarioles expressing GFP-Piwi (greyscale) in the indicated genotypes (captions to the right show enlarged germaria). (E) Top: Confocal images (scale bars: 50 μm) showing ovarioles stained for Aubergine in the indicated genotypes (captions to the right show enlarged germaria). Bottom: Western blot analysis indicating levels of endogenous Aubergine in ovarian lysates from flies with the indicated genotypes (anti ATP-synthase blot served as loading control).

To evaluate the efficiency of the different drivers in inducing transgenic RNAi, we crossed them with flies carrying a very potent UAS-shRNA[GFP] transgene and one CRISPR-modified *piwi* allele harboring an N-terminal GFP-tag. As expected, MTD-Gal4 and *tj*-Gal4 induced strong depletion of GFP-Piwi in the entire ovarian germline or soma, respectively (Figure 2D). For *osk*-Gal4 and *αTub67C*-Gal4, GFP-Piwi levels were reduced from stage 2 egg chambers onwards and were undetectable in older egg chambers. As a more quantitative assay, we crossed the different Gal4 drivers with a UAS-shRNA[*piwi*] line and determined female sterility. shRNA-mediated depletion of Piwi with MTD-Gal4 resulted in 100% sterility (n = 200 laid eggs). For *osk*-Gal4 or *αTub67C*-Gal4, we observed near-complete sterility with occasional escapers. A driver line combining *αTub67C*-Gal4 and *osk*-Gal4 on the second chromosome, henceforth designated as TOsk-Gal4, resulted in complete sterility and was therefore used throughout this study.

We compared TOsk-Gal4 and MTD-Gal4 in the context of the germline piRNA pathway and depleted the two central Argonaute proteins Piwi or Aubergine (Aub) with UAS-shRNA lines. Endogenous Piwi or Aub proteins were undetectable in all germline cells for MTD-Gal4 and from stage 2/3 egg chambers onwards for Tosk-Gal4 (shown for Aub in Figure 2E). For both Gal4 drivers, depletion of Piwi or Aub resulted in complete female sterility. To compare how depletion of Piwi with MTD-Gal4 versus TOsk-Gal4 impacts TE silencing, the central function of the germline piRNA pathway in *Drosophila*, we conducted RNA-seq experiments and, for selected TEs, RNA fluorescent in situ hybridization (FISH) experiments on ovaries with the different knockdown conditions. Overall, the same TEs that were de-repressed in ovaries depleted for Piwi during the entirety of oogenesis (MTD-Gal4) were also de-repressed, albeit at lower levels, in ovaries where transgenic RNAi was effective only from stage 3 egg chambers onwards (TOsk-Gal4) (Figure 3A). Examples for TEs exhibiting similar de-repression behavior are *blood* or *HMS Beagle* (Figures 3B). Mid and late stage egg chambers (where loss of Piwi is indistinguishable in MTD-versus TOsk-Gal4 crosses) contribute the bulk of the ovary mass and RNA. We therefore argue that the milder TE de-repression in the TOsk-Gal4 crosses is not due to differences in knockdown efficiency, but rather due to delayed TE de-silencing upon loss of piRNA pathway activity. Piwi-mediated heterochromatin formation at TE loci likely contributes to this pattern. In agreement with this, steady state RNA levels for the LTR retrotransposon *mdg3*, which is primarily repressed via Piwi-mediated heterochromatin formation (Senti *et al*. 2015) and whose steady state RNA levels were more than 350-fold elevated upon MTD-Gal4-mediated Piwi knockdown, did not change more than ∼2-fold when Piwi was depleted with Tosk-Gal4 (Figures 2A, C). Taken together, TOsk-Gal4 allows potent transgenic RNAi in germline cells of maturing egg chambers. In the case of the piRNA pathway, this results in a temporal delay in TE de-repression compared to a pan-germline knockdown.

**Figure 3:**
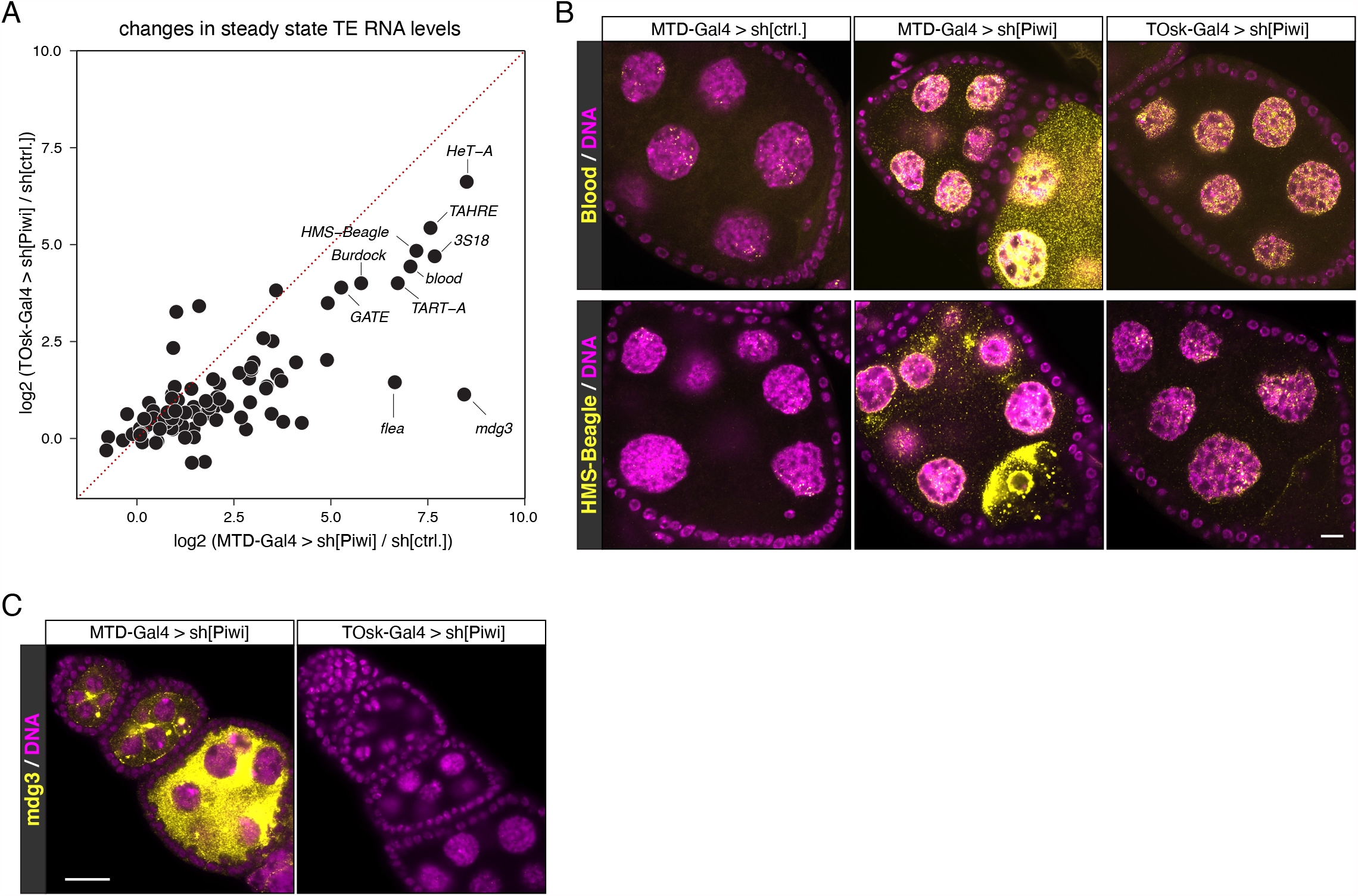
Comparison of MTD-Gal4 and TOsk-Gal4 driven transgenic RNAi. (A) Scatter plot showing log2 fold changes (respective to control) of TE steady state RNA levels in ovaries where germline Piwi was depleted using MTD-Gal4 or TOsk-Gal4. (B and C) Confocal images (scale bars: 10 μm) of egg chambers with indicated genotype stained for the TEs *blood* or *HMS-Beagle* (B), or for *mdg3* (C) using RNA-FISH (yellow; DAPI: magenta).

### Intersection points between piRNA pathway and essential cellular processes

To evaluate the utility of the TOsk-Gal4 transgenic RNAi system, we investigated biological processes that are required for transposon silencing and for cellular viability. We focused on the nuclear RNA export factors UAP56 and Nxf1, the protein exporter Crm1 (Zhang *et al*. 2012; ElMaghraby *et al*. 2019; Kneuss *et al*. 2019), and on the protein SUMOylation machinery with the E1 activating enzyme Uba2–Aos1 and the E3 Ligase Su(var)2-10 (Ninova *et al*. 2020). Depletion of any of these factors with MTD-Gal4 resulted in rudimentary ovaries, precluding any meaningful analysis as these lacked germline tissue, evidenced by the absence of Aub expressing cells (Figures 4, 5A; shown for UAP56, Crm1, Sbr). Crossing the same set of UAS-shRNA lines with the Tosk-Gal4 driver yielded flies with partially restored ovarian morphology and germline development.

**Figure 4:**
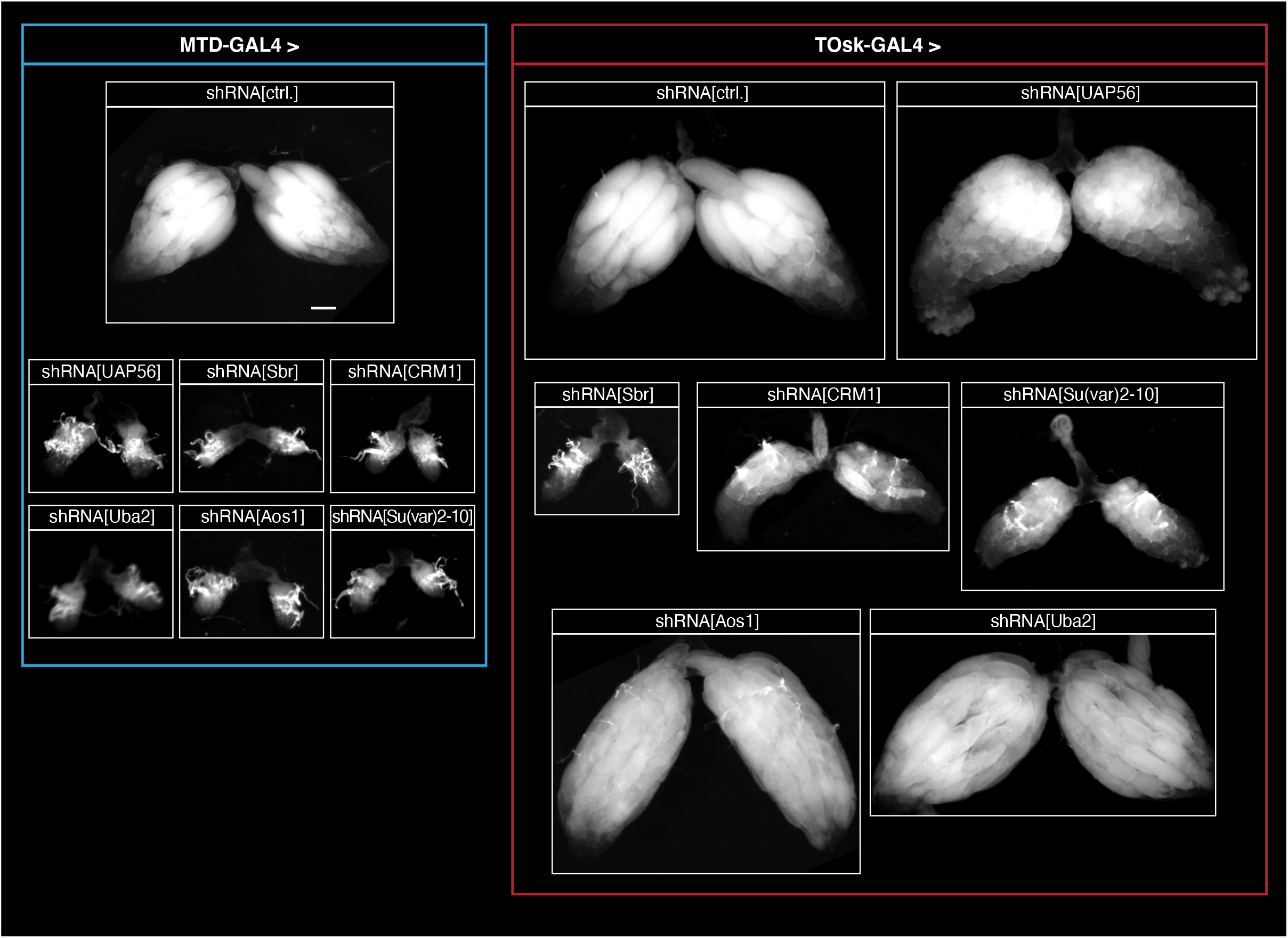
Transgenic RNAi of essential genes with TOsk-Gal4. Bright field images (scale bar for all images: 200 μm) showing ovarian morphology from flies of the indicated genotype (to the left: MTD-Gal4 crosses; to the right: TOsk-Gal4 crosses).

**Figure 5:**
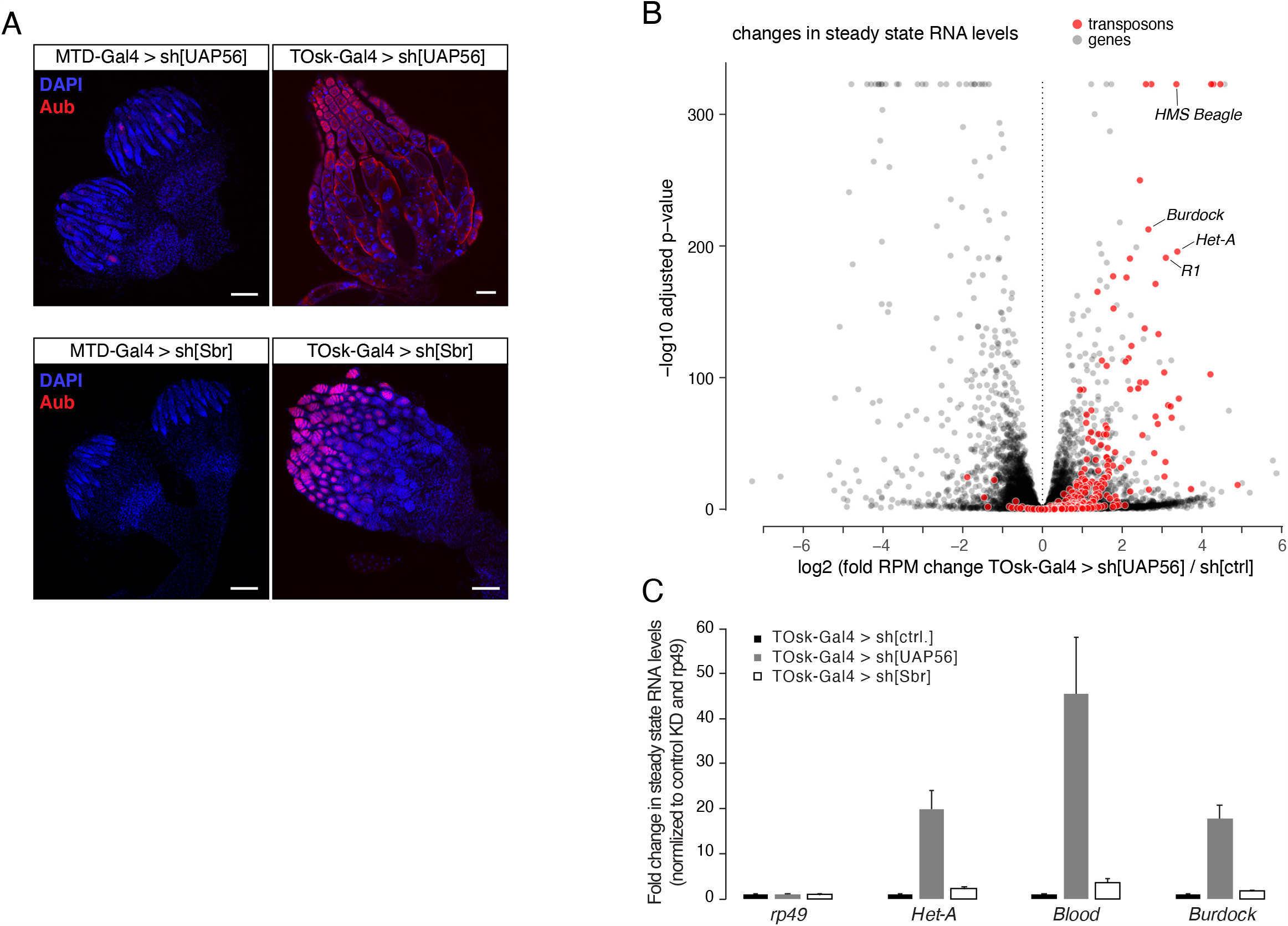
Dual role of UAP56 in mRNA export and transposon defense. (A) Confocal images (scale bars: 100 μm) showing whole ovaries from flies of indicated genotypes stained with anti-Aubergine antibody (red); DNA stained with DAPI (blue). (B) Volcano plot showing fold changes in steady state mRNA (black dots) and TE transcript levels (red dots) in ovaries from TOsk-Gal4 > sh[UAP56] flies versus control flies (n = 2 biological replicates). (C) qRT-PCR analysis showing fold changes in steady state TE transcript levels in ovaries from flies with indicated genotype qRT-PCR (n = 2 biological replicates; normalized to *rp49* mRNA levels).

Nuclear export of mRNA and piRNA precursors, both transcribed by RNA Polymerase II, requires the THO complex and the RNA Helicase UAP56 (Zhang *et al*. 2012; Hur *et al*. 2016; Zhang *et al*. 2018; ElMaghraby *et al*. 2019). Together, these proteins license the loading of the RNA cargo onto a specific nuclear export receptor belonging to the NXF protein family (Nxf1 for mRNA, Nxf3 for piRNA precursors), which subsequently shuttles its respective RNA cargo through nuclear pore complexes into the cytoplasm (Kohler and Hurt 2007; ElMaghraby *et al*. 2019; Kneuss *et al*. 2019). Consistent with their central role in nuclear mRNA export, RNAi-mediated depletion of Nxf1 (*Drosophila* Small bristles; Sbr) or UAP56 with MTD-Gal4 resulted in ovaries lacking germline cells (absence of Aub expressing cells; Figure 5A). Depletion of UAP56 or Sbr with TOsk-Gal4 also yielded sterile females. These flies, however, contained larger ovaries with clearly developing egg chambers, hence permitting molecular analyses (Figures 4, 5A). We performed RNA-seq experiments on ovaries where UAP56 was depleted with TOsk-Gal4. Besides many genes that were de-regulated compared to control ovaries, numerous piRNA pathway repressed TEs were de-silenced (Figure 5B). In contrast, TEs were not de-repressed in ovaries depleted of the essential mRNA export receptor Sbr (Figure 5C) supporting a direct role of UAP56 in the piRNA pathway, beyond nuclear export of mRNAs encoding for piRNA pathway proteins. These findings highlight the dual role of UAP56 as a central gate keeper to feed RNA cargo into two distinct nuclear export receptors, Nxf1–Nxt1 for mRNAs and Nxf3–Nxt1 for Rhino dependent piRNA precursors.

As a second intersection point between piRNA pathway and essential cellular processes, we chose piRNA-guided heterochromatin formation. The nuclear Argonaute protein Piwi, guided by its bound piRNAs, induces transcriptional gene silencing and specifies the local formation of heterochromatin at genomic TE insertions (Wang and Elgin 2011; Sienski *et al*. 2012; Le Thomas *et al*. 2013; Rozhkov *et al*. 2013). This process depends on transcription of a piRNA-complementary nascent transcript. To mediate silencing, piRNA-loaded Piwi requires a handful of piRNA pathway-specific factors (Gtsf1/Asterix, Maelstrom, SFiNX complex) as well as factors of the general heterochromatin machinery that the piRNA pathway taps into (Czech *et al*. 2018; Ninova *et al*. 2019). Depletion of these general factors via MTD-Gal4 driven transgenic RNAi yields rudimentary ovaries with absent germline tissue. To explore the utility of the TOsk-Gal4 system, we focused on the protein SUMOylation pathway that is involved in numerous chromatin-related processes and that is required for Piwi-mediated transcriptional silencing and heterochromatin formation (Gareau and Lima 2010; Jentsch and Psakhye 2013; Ninova *et al*. 2020).

Protein SUMOylation requires E1 and E2 enzymes. *Drosophila* expresses a single E1 enzyme (Aos1– Uba2) and a single E2 enzyme (Lwr), which in a stepwise manner transfer a SUMO moiety onto a target Lysin of the substrate. A handful of E3 ligases potentiate the SUMOylation process in a substrate specific manner. Recent work has uncovered a critical role for the E3 ligase Su(var)2-10, and hence SUMOylation, in Piwi-mediated heterochromatin formation (Ninova *et al*. 2020). Depletion of Uba2, Aos1 or Su(var)2-10 with MTD-Gal4 was incompatible with GSC survival and oogenesis (Figure 4) (Yan *et al*. 2014). We therefore used TOsk-Gal4 driven transgenic RNAi to probe for a requirement for SUMOylation in the piRNA pathway. Piwi-mediated transcriptional silencing and heterochromatin formation can be mimicked by experimental tethering of the SFiNX complex to a nascent transcript using the λN-boxB system (Figure 6A) (Sienski *et al*. 2015; Yu *et al*. 2015). In ovaries from flies that ubiquitously express a GFP reporter with boxB sites and that harbor the TOsk-Gal4 driver and a UASp-λN-Panoramix (SFiNX subunit) construct, GFP expression was silenced specifically in the germline from stage 3 egg chambers onwards (Figure 4B). Simultaneous expression of shRNA constructs targeting Piwi, which acts upstream of SFiNX, had no impact on GFP silencing. Similarly, targeting the mRNA exporter Sbr (Nxf1) did not interfere with SFiNX function, supporting the specificity of the assay. In contrast, depletion of the SUMOylation machinery (Aos1, Uba2 or Su(var)2-10) restored GFP expression confirming that protein SUMOylation is required for SFiNX-mediated heterochromatin formation (Figure 6B).

**Figure 6:**
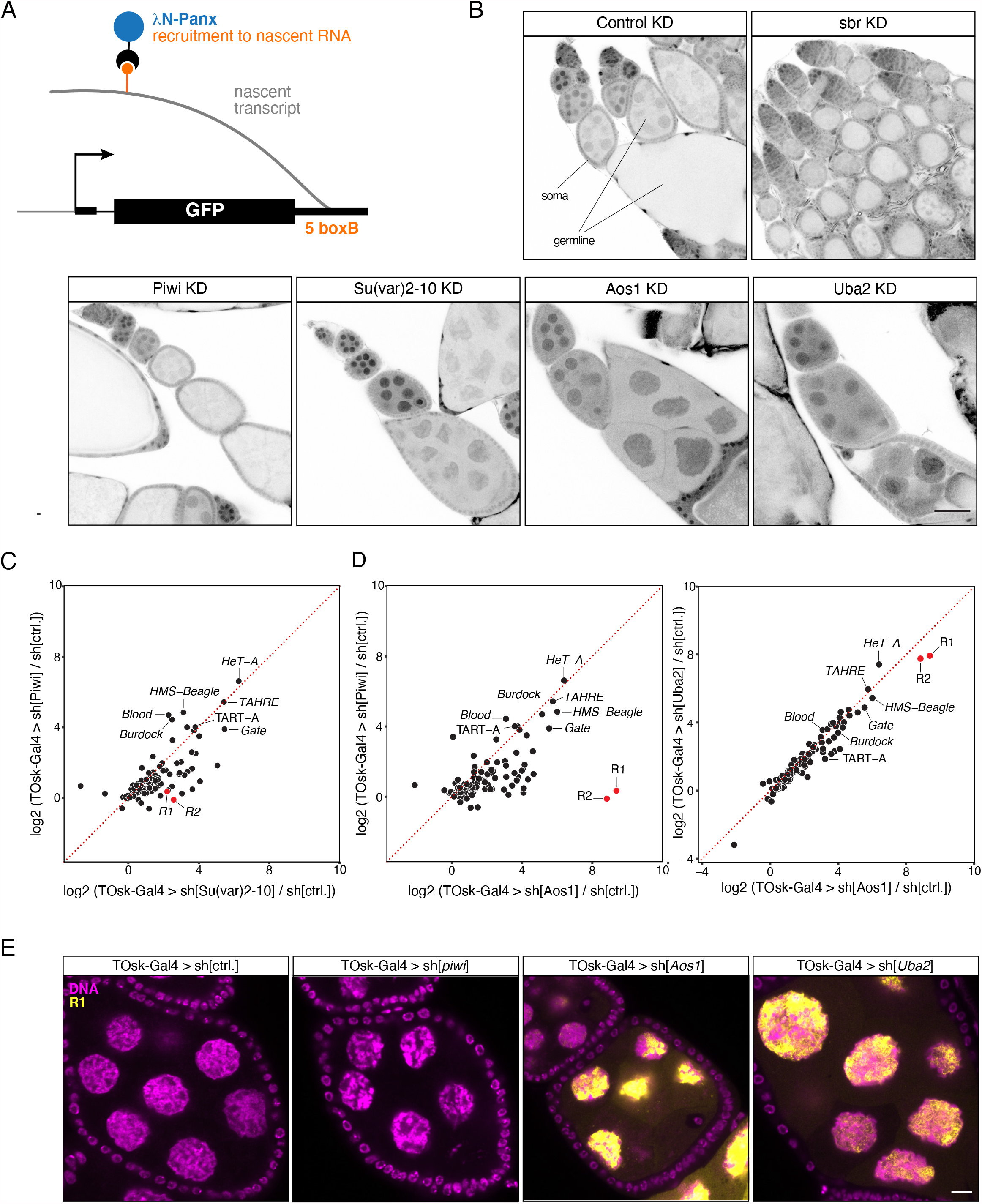
The SUMO machinery is required for TE repression in the germline. (A) Cartoon depicting the transgenic RNA-tethering reporter based on the λN-boxB system. The α- tubulin promoter expresses GFP in all cells, and the 3′ UTR harbors five boxB sites to allow tethering of λN-Panx to the reporter mRNA. (B) Confocal images (scale bar: 50 μm) showing GFP signal (greyscale) in egg chambers expressing the GFP-boxB reporter plus λN-Panx and the indicated shRNAs specifically in the germline using TOsk-Gal4 (somatic cells serve as control). (C-D) Scatter plots based on RNA-seq data showing log2 fold changes (relative to control) in TE steady state transcript levels in ovaries from flies of the indicated genotype. (E) Confocal images (scale bar: 10 μm) of egg chambers from flies with indicated genotype showing *R1* transposon mRNA using RNA-FISH (yellow; DAPI: magenta).

To obtain a more quantitative and systematic impact of the SUMOylation pathway on TE silencing in the female germline, we performed RNA-seq experiments on ovaries depleted (via TOsk-Gal4) for Piwi, Uba2, Aos1 or Su(var)2-10 and compared TE RNA levels to those in control ovaries. Depletion of Piwi or Su(var)2-10 resulted in overall similar TE silencing defects (Figure 6C). When comparing TE transcript levels in Piwi depleted ovaries to those in ovaries depleted for the SUMO E1-ligase subunits Aos1 or Uba2, a similar set of TEs showed increased expression (Figure 6D). However, the *R1* and *R2* retrotransposons, two LINE elements that integrate specifically into rDNA units, were strong outliers as they showed nearly exclusive de-repression in ovaries lacking Aos1 or Uba2. In Aos1 or Uba2 depleted ovaries, steady state RNA levels of *R1* and *R2* reached enormous levels and were among the most abundant cellular transcripts (Figures 6D, E). *R1* or *R2* showed no de-repression in ovaries lacking Piwi (even when depleted via the MTD-Gal4 driver) although germline Piwi is loaded with *R1* and *R2* derived piRNAs. In agreement with a recent report (Luo *et al*. 2020), our data indicate that the SUMOylation pathway is integral for silencing *R1* and *R2*, likely in a piRNA-independent and to a large extent also in a Su(var)2-10 independent manner.

### A TOsk-Gal4 system for long dsRNA hairpins and earlier expression

During our studies on the transgenic RNAi system using TOsk-Gal4, we encountered two technical aspects that warranted further modifications. While TOsk-Gal4 was highly efficient in depleting target genes using shRNA lines (e.g. TRIP collection), it was very inefficient with long dsRNA hairpin lines (VDRC collection). For example, when we crossed TOsk-Gal4 to UAS lines harboring long dsRNA hairpins targeting the SUMO-pathway, the resulting females were fertile, in stark contrast to crosses to shRNA lines targeting the same genes. This was reminiscent of the pan-oogenesis Gal4 driver system where efficient transgenic RNAi using long hairpin constructs requires the co-expression of the siRNA generating ribonuclease Dcr-2 (Wang and Elgin 2011). Indeed, when we combined TOsk-Gal4 with an X-chromosomal UAS-Dcr-2 transgene, knockdown of Smt3 (*Drosophila* SUMO) as well as of the single E2 SUMO-conjugating enzyme Ubc9 (encoded by the *lwr* gene) yielded sterile females. These exhibited severe reductions in Smt3 levels specifically in the germline (for the *smt3* knockdown) and the characteristic strong de-repression of the *R1* and *R2* elements (Figure 6A, B). Given the almost genome wide collection of dsRNA hairpin lines at the VDRC, the TOsk-Gal4 > UAS-Dcr-2 combination stock will enable systematic genetic screens targeting genes that with a pan-oogenesis knockdown would yield rudimentary ovaries, often lacking germline tissue.

We finally considered the timing of oogenesis in respect to the onset of transgenic RNAi. The developmental process from germline stem cell division to mature egg takes nearly one week (Horne-Badovinac and Bilder 2005; He *et al*. 2011). Up to four days are spent during the germarium stages, meaning before the onset of measurable depletion of target proteins using TOsk-Gal4. Though extraordinary efficient in depleting even abundant factors like Piwi or Smt3, this means that the time window of efficient transgenic RNAi is around two to three days. To extend this effective knockdown period, we turned to the *bam*-Gal4 driver, which is expressed in a narrow time period of around one day at the onset of cystoblast division (Figure 7C) (Chen and McKearin 2003). When combining *bam*-Gal4 with TOsk-Gal4 (BamTOsk-Gal4), the knockdown of GFP-Piwi with an shRNA line against GFP indicated severe loss of Piwi already at the germarium stage 2b, thereby extending the entire knockdown window of this triple driver to three to four days (Figure 7D). For a direct comparison of the various Gal4 driver combinations, we used the sh[*sbr*] line that leads to a highly potent depletion of the cell-essential mRNA export receptor Sbr (*Drosophila* Nxf1) (Figure 7E). Depletion of Sbr with the pan-oogenesis MTD-Gal4 driver resulted in the complete absence of germline tissue (no Aub positive cells). Depletion by TOsk-Gal4 resulted in phenotypically normal germaria and two to four additional egg chambers per ovariole. The BamTOsk-Gal4 cross resulted in an intermediate phenotype with normal germaria but only one additional egg chamber per ovariole. Thus, the BamTOsk-Gal4 driver represents an ideal driver to study gene function in the differentiating female germline via transgenic RNAi.

**Figure 7:**
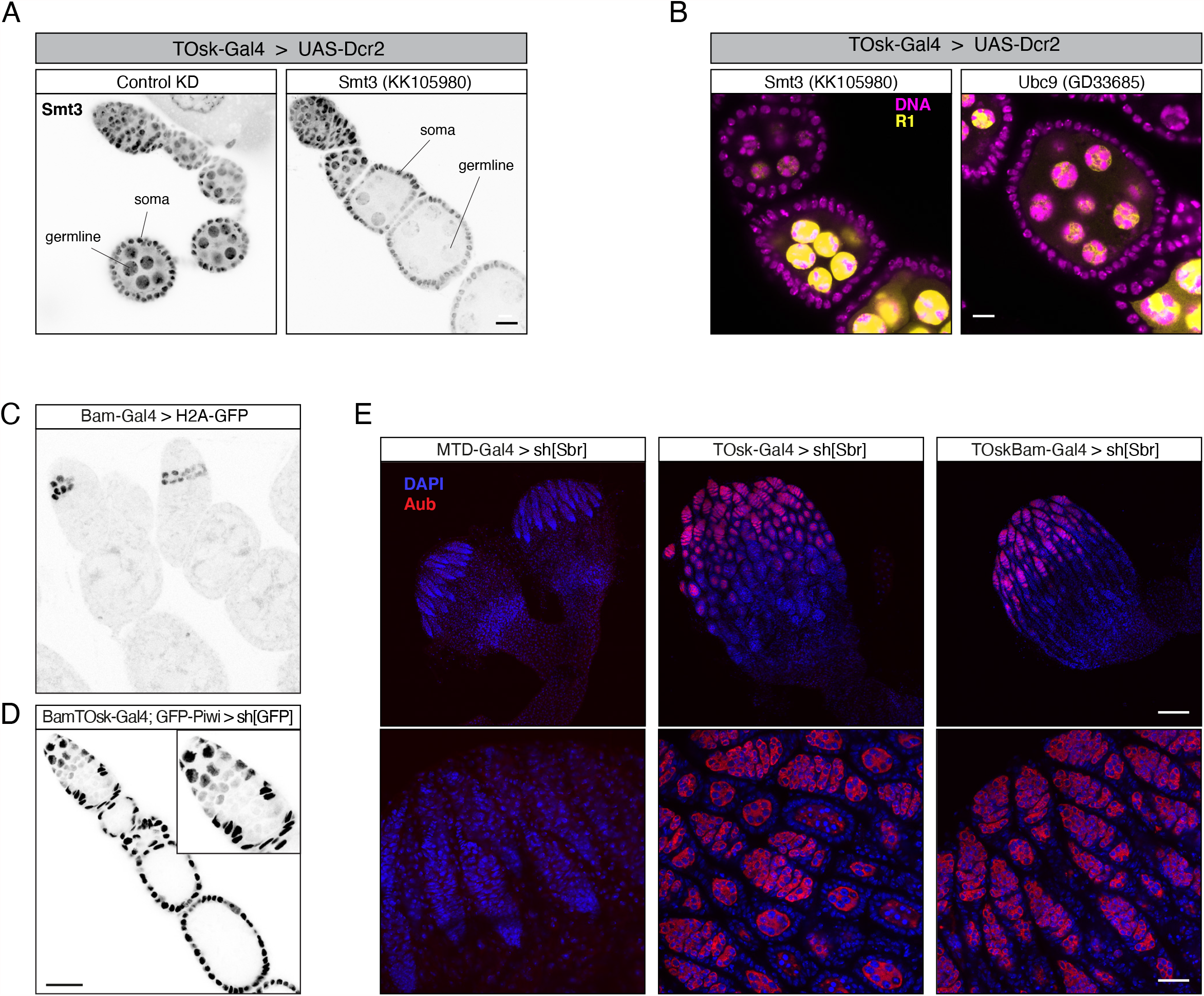
Extensions of the TOsk-Gal4 system. (A) Confocal images (scale bar: 10 μm) showing ovarioles from flies with indicated genotype stained with anti-Smt3 antibody (greyscale). (B) Confocal images (scale bar: 10 μm) of egg chambers from flies with indicated genotype showing *R1* transcripts (RNA-FISH: yellow, DAPI: magenta). (C) Confocal image showing ovarioles expressing H2A-GFP driven by the *bam*-Gal4 driver. (D) Confocal image (scale bar: 25 μm) showing GFP-Piwi levels (greyscale) in an ovariole expressing an sh[GFP] transgene with BamTOsk-Gal4 (inset shows the enlarger germarium; somatic cells serve as internal control). (D) Confocal images showing whole ovaries (top row; scale bar: 100 μm) from flies of indicated genotype stained with anti-Aubergine antibody (red); DNA stained with DAPI (blue). Early oogenesis regions are highlighted in the bottom row (scale bar: 25 μm).

Taken all together, our work provides a versatile, highly specific and powerful genetic toolkit that permits tissue specific RNAi at various stages of oogenesis in conjunction with GFP markers for the visualization of subcellular structures. While our focus was on piRNA biology, the presented approach is applicable to any expressed gene in the ovarian germline and complements previously established assays based on MTD-Gal4 or a double *αTub67C*-Gal4 driver (Staller *et al*. 2013; Yan *et al*. 2014). Through the compatibility with genome wide short and long UAS-dsRNA lines available from the Bloomington stock center or the VDRC, our work enables reverse genetic screens for the involvement of cell-essential factors in specific biological processes.

## ACKNOWLEDGEMENTS

We thank D. Handler and M. Gehre for help with bioinformatics and computational analyses, the IMBA/IMP/GMI core facilities, in particular BioOptics for support, the Vienna Biocenter Core Facilities (VBCF) for providing NGS, COVID-19 testing, and fly stocks and husbandry services (VDRC). Imre Gaspar and Daniel St. Johnston provided fly stocks. We sincerely thank Clemens Plaschka and Brennecke lab members for support and insightful discussions.

## AUTHOR CONTRIBUTIONS

The project was conceived by JB, MFE and LT. MFE and LT performed all molecular biology and genetics experiments. KAS conceptualized the BamTOsk-Gal4 driver and KAS and KM generated essential reagents and fly stocks. JB supervised the study and MFE, LT and JB wrote the paper with input from all authors.

## FIGURE LEGENDS

**Figure S1:**
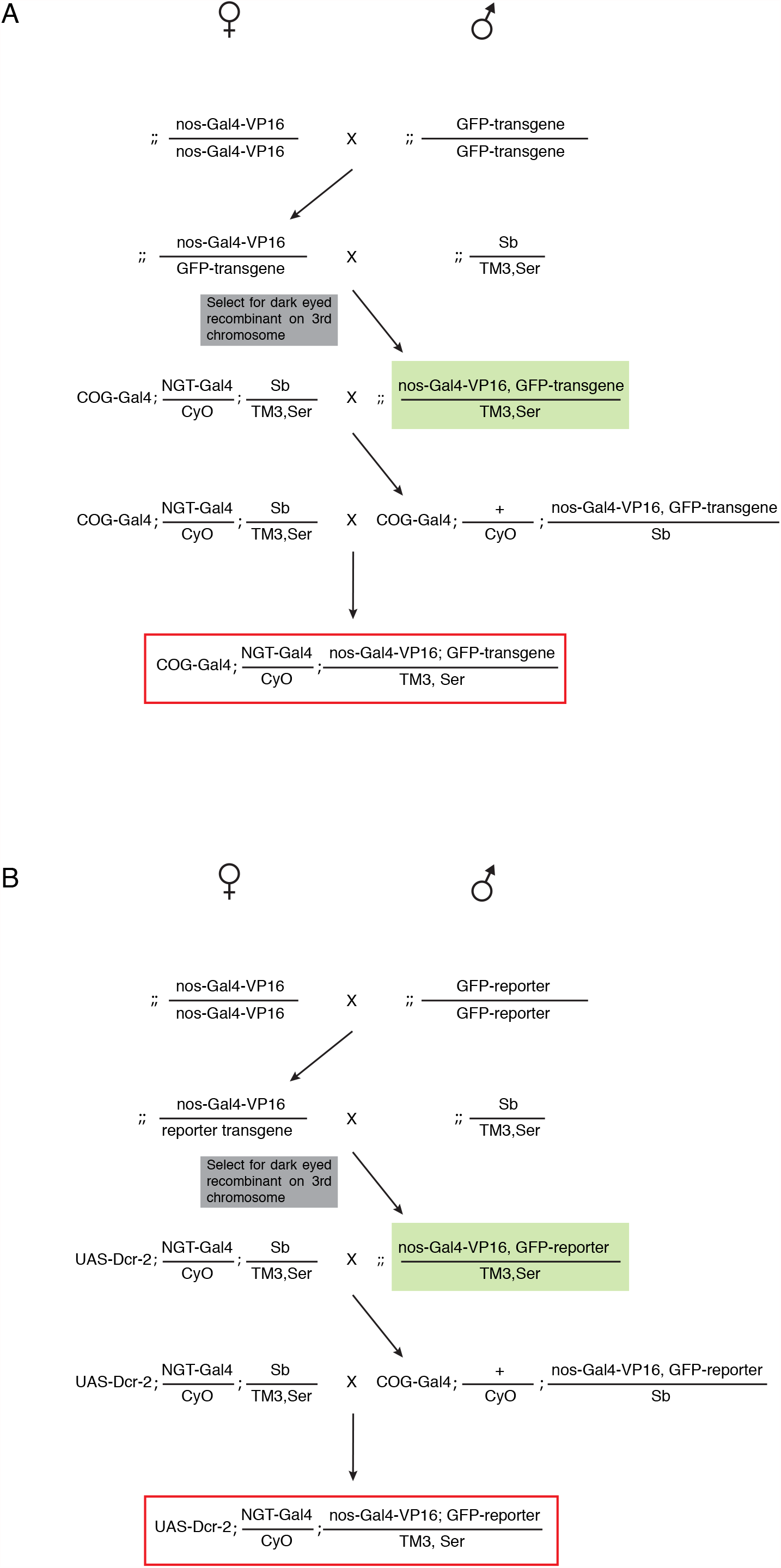
Crossing scheme to generate marker lines with Gal4 Drivers. (A) Scheme for construction of MTD-Gal4 lines with compatible with short hairpin RNA (shRNA) UAS-lines. (B) Scheme for construction of nanos-Gal4 lines with a UAS-Dcr-2 transgene compatible with long hairpin RNA UAS-lines.

